# Design and synthesis of nanocarriers containing posaconazole against etiological agents of mucormycosis

**DOI:** 10.1101/2025.05.08.652746

**Authors:** Parsa Panahi, Omid Pourdakan, Hannaneh Zarrinnahad, Shahram Mahmoudi, Ghazal Aghaie, Farnaz Hosseini, Sayyed Amirpooya Alemzadeh, Mohammadamin Malek Raeesi, Hossein Mirhendi, Bita Mehravi

## Abstract

**Background:** Posaconazole is an effective antifungal agent used for many fungal diseases, and its application has grown recently. Posaconazole was studied as a treatment option after the increase in mucormycosis cases linked to COVID-19, and promising results were obtained. However, despite its effectiveness, researchers are looking into new approaches to drug delivery due to problems including restricted bioavailability and short half-life. To address these issues, this project intends to create and synthesize posaconazole-containing liposomes.

**Methods:** In this study, according to the synthesis of Posaconazole-loaded liposomes, their physical and chemical properties were assessed using SEM, DLS, and FTIR techniques. The drug loading capacity in the nanoparticles, its release in vitro, and the antifungal activity against mucormycosis-causing fungi were evaluated. Finally, the cytotoxicity of these nanoparticles was assessed using the MTT Assay.

**Results:** The synthesized nanoparticles were spherical with an average size of approximately 60 nm. The drug loading efficiency was found to be 78.41%. The results indicated that the posaconazole nanoparticles demonstrated a slow and continuous drug release over five days. These nanoparticles demonstrated higher efficacy than posaconazole individually in antifungal tests. Additionally, cytotoxicity tests revealed that, up to a 400 µg/ml concentration, posaconazole-loaded nanoparticles were less toxic than free posaconazole.

**Conclusions:** These nanoparticles may provide a useful therapeutic option for the treatment of mucormycosis while circumventing the disadvantages of posaconazole due to their improved antifungal activities and decreased toxicity compared to free posaconazole.

## Introduction

Mucormycosis is a class of invasive fungal infections that are historically related to zygomycetes and are recognized based on their characteristic ribbon-like appearance. (1). Given recent changes in taxonomy, those organisms have been divided into two main orders: Entomophthorales and Mucorales ((2), (3)).

Mucormycosis, commonly known as “black fungus,” refers to acute or subacute infections caused by fungi in the Mucorales order, recognized for their high growth rates. This condition predominantly affects immunocompromised individuals, particularly those suffering from uncontrolled diabetes mellitus, and is associated with severe complications, including progressive tissue necrosis and high mortality rates ((1),(4)).

Although rare cases of mucormycosis have also been documented in individuals with intact immune systems (5). Mucormycosis reveals different clinical manifestations that affect various anatomical sites including the skin and kidneys central nervous system and gastrointestinal tract.

The most prevalent and severe forms include rhino-orbit-cerebral and pulmonary mucormycosis. In addition, isolated infections have been documented affecting the heart (and its valves), uterus, bladder, lymph nodes, mediastinum, middle ear, parotid gland, and other organs. (6).

This type usually develops in immunocompromised patients after inhalation of Rhizopus spores into the sinuses, where the spores germinate rapidly and develop into numerous hyphae. Vascular and neurovascular spread occurs rapidly through the infiltration of the vascular wall (7).

Thrombosis and neurological impairment can occur due to direct tissue invasion. Additionally, the involvement of blood vessels, bone, cartilage, nerves, perineural spaces, and meninges is frequently observed. Tissue necrosis in the palate may lead to erosion of the palatine bone and destruction of the turbinates. The infection continues to spread toward the orbital cavity, eventually penetrating the brain, where it causes cranial nerve damage, widespread blood clotting, and blockage of the carotid artery. The cumulative effects of the infection result in severe tissue destruction in the head and facial regions, often leading to death in immunocompromised patients (8, 9).

The efficient management of mucormycosis necessitates early detection of the fungal infection. Amphotericin B is the primary antifungal agent used to treat this condition. The liposomal formulation of this drug is administered intravenously at high doses once daily as initial therapy. The duration of amphotericin B treatment is contingent upon the patient’s clinical status. To transition to oral posaconazole or isavuconazole, intravenous amphotericin B is gradually tapered after several weeks of initial therapy and subsequent clinical improvement. It is important to note that oral posaconazole suspensions are not recommended due to their poor bioavailability and inadequate gastrointestinal absorption. For patients who exhibit intolerance or do not respond to amphotericin B, oral posaconazole may be considered. Numerous studies have established the efficacy of posaconazole in treating mucormycosis, particularly in cases involving the rhino-orbit-cerebral form (13, 14, 15). However, challenges arise in connection with the use of posaconazole. Monitoring serum concentrations one week after the initiation of therapy is crucial to ensure that levels remain above 1 microgram per milliliter, which is considered the effective concentration, due to its low bioavailability. Consequently, researchers and clinicians are investigating topical posaconazole as a potential treatment for orbital mucormycosis (6). Novel drug delivery methods have emerged as promising strategies to enhance the effectiveness of pharmaceuticals. Liposomal drug delivery is one such method that has been extensively studied for various applications, including gene delivery and drug formulation. Liposomes are spherical vesicles that encapsulate aqueous material within a lipid bilayer, which may consist of phospholipids or synthetic amphiphiles. Liposomal encapsulation reduces toxicity and alters pharmacokinetics and pharmacodynamics, thereby improving therapeutic efficacy. Both hydrophobic and hydrophilic drugs can be encapsulated within liposomes; hydrophobic drugs are associated with the lipid bilayer, while hydrophilic drugs are entrapped within the aqueous core.

Our goal is to enhance the therapeutic efficacy of posaconazole by encapsulating it in liposomes; this approach reduces the frequency of doses and minimizes the necessity for multiple injections in patients with rhino- orbit-cerebral mucormycosis by reducing the dose frequency.

## Material and Methods

### Materials

#### Synthesis of liposomes containing the drug posaconazole

The synthesis of liposomes containing the drug posaconazole was performed using the thin film method (19). Initially, 10 mg of posaconazole was weighed and combined with 300 µl of dichloromethane (DCM), and the mixture was stirred for 10 minutes to ensure complete dissolution. Concurrently, 100 µl of lipid solution (phosphatidylcholine) was mixed with 200 µl of DCM in a separate container and stirred for 10 minutes until the lipid was fully dissolved. Once both the drug and lipid solutions were completely dissolved in DCM, the drug solution was added to the lipid solution and stirred for an additional 30 minutes at room temperature. Eventually, the mixture containing lipid and posaconazole was placed in a rotary evaporator for 60 minutes at 60°C under reduced pressure (100 mbar) to evaporate the DCM, forming posaconazole liposomes. The resulting film was then dissolved in 20 ml of distilled water and stirred for 30 minutes to achieve homogeneity. The solution was placed in an ice bath and subjected to probe sonication for 10 minutes at a power of 200 watts with cycles of 10 seconds on and 3 seconds off to minimize liposome size. After sonication, the final solution was transferred to microtubes and centrifuged at 18,000 rpm for two rounds of 15 minutes each. The posaconazole liposomes settled at the bottom of the microtubes, and the supernatant was removed for later analysis of free drug content to calculate drug loading efficiency. Finally, the microtubes containing the liposome precipitate were stored at -80°C for 24 hours before being placed in a freeze-dryer to obtain a dry powder of posaconazole liposomes for subsequent experiments. To determine the maximum absorption wavelength of posaconazole, 1 mg of the drug was dissolved in 2 ml of DCM to create a stock solution from which concentrations ranging from 0 to 40 µg/ml were prepared. UV spectrophotometer measurements revealed that 260 nm was the maximum wavelength of absorption. Excel software was used to develop an absorbance versus concentration calibration curve for each prepared solution at this wavelength. It is noteworthy that DCM was used as a blank in each measurement step. An electron microscope image was obtained by using a sputter coater to coat powdered samples with gold .The samples were placed in a vacuum chamber where argon gas replaced the air, ionizing argon atoms that deposited gold onto the sample surface. As a result of this preparation, scanning electron microscopy (SEM) was used to evaluate the particle morphology and size distribution of the samples. The particle size of powdered samples was assessed by dynamic light scattering (DLS) after ultrasonic treatment using powdered samples in distilled water. For evaluating nanoparticle stability, zeta potential measurements were made. Stable particles are those with zeta potentials greater than 30 mV or less than -30 mV. Infrared spectroscopy (FTIR) analysis involved mixing one gram of powdered sample with KBr at a ratio of 1:100 and pressing it into a pellet suitable for infrared examination over a range of 400-4000 cm1. The drug loading capacity within albumin-coated liposomes was determined by analyzing the supernatant from centrifuged solutions containing unencapsulated drugs. The amount of unencapsulated drug was calculated based on absorbance measurements at the maximum wavelength. The percentage drug loading was calculated using the formula:

Loading=100×(Total Drug−Free Drug)Total DrugLoading=Total Drug100×( Total Drug−Free Drug)

#### Assessment of Drug Release from Liposomes

To study drug release kinetics, dialysis bags containing nanoparticles were immersed in phosphate-buffered saline (PBS). The PBS buffer was prepared by mixing sodium chloride, potassium chloride, disodium phosphate, and monopotassium phosphate to achieve a final volume of one liter. After activating dialysis bags by boiling them in distilled water, nanoparticles containing posaconazole were dispersed in PBS and placed within these bags. At predetermined intervals (1 hour, 2 hours, etc.), samples were withdrawn from the external PBS solution and replaced with fresh buffer. A standard curve was used to quantify released drug concentrations using UV spectrophotometry. Six clinical isolates identified through ITS mapping have been tested for efficacy with posaconazole liposomes against mucormycosis-causing fungi. Initial fungal cultures were established on Sabouraud dextrose agar plates prepared under sterile conditions. For antifungal activity assessment, RPMI-1640 medium without bicarbonate was prepared according to Clinical & Laboratory Standards Institute (CLSI) guidelines. Each isolate was tested with posaconazole stock solutions (16 g/mL) along with fungal suspensions. In 96-well plates, serial dilutions of each isolate at controlled temperatures were performed to evaluate antifungal activity. The viability of L929 fibroblasts was assessed using F12 DMEM supplemented with fetal bovine serum and antibiotics. MTT assays were performed following exposure to different concentrations of posaconazole and its liposomal formulation. To assess the safety profile of posaconazole-loaded liposomes in this study, cytotoxicity studies on mammalian cells were conducted to determine their synthesis, characterization, and evaluation processes.

### Characterization of Nanoparticles

#### Determination of the Maximum Absorption Wavelength of Posaconazole

To determine the maximum absorption wavelength and calibration curve of the drug posaconazole, 1 mg of the drug was initially dissolved in 2 ml of its solvent (Dichloromethane, DCM) to prepare a stock solution. From this stock solution, further dilutions were made to achieve concentrations ranging from 0 to 40 µg/ml of posaconazole. Subsequently, using a UV spectrophotometer, the wavelengths from 200 to 500 nm were examined, leading to the identification of the drug’s maximum absorption wavelength at 260 nm. The absorbance of each solution at the specified concentrations was then measured at this wavelength, allowing for the construction of an absorbance versus concentration (in µg/ml) graph. Data analysis was performed using Excel software. It is noteworthy that the DCM solvent was utilized for blanking the instrument at each measurement stage.

#### Sample Preparation for Examination with SEM

For imaging powder samples using a Scanning Electron Microscope (SEM), the samples were initially coated with a thin layer of gold using a Sputter Coater. Once the samples were placed in a dedicated chamber, the air within the chamber was evacuated and replaced with argon gas (inert). An electrical potential difference was established between the cathode (gold disk) and anode, causing argon to be ionized and atoms to move from the gold surface to the sample surface. Subsequently, a metallic layer approximately 20 to 30 nm thick was deposited onto the sample. Following electron beam irradiation, the gold atoms released electrons and began the imaging process. The preparation of samples for SEM is essential for ensuring high-quality imaging as well as accurate composition and morphology analysis.

#### Sample Preparation for Analysis with DLS

A probe sonicator was used to disperse powder samples in 1 ml of distilled water using Dynamic Light Scattering (DLS) to ensure uniform dispersion and prepare them for DLS analysis. Additionally, to assess the stability of the nanoparticles in this study, the zeta potential of the samples was also evaluated. Nanoparticles exhibit optimal stability when their surface charge is greater than +30 mv or less than -30 mv.

#### Functional Group Analysis via FTIR Method

For this test, 1 mg of the powdered solid sample was thoroughly mixed with 100 mg of dry potassium bromide (KBr) in a weight ratio of 1:100. The mixture was then placed in a specialized metal mold and subjected to pressure using a hydraulic press to form a transparent pellet. This process is based on the principle that halide salts, when subjected to sufficient pressure, can be transformed into a glassy pellet that is transparent to infrared radiation. This property is utilized for preparing solid samples for analysis. Following the preparation of the pellet, analysis was conducted using an FTIR spectrometer over the wavenumber range of 4000 to 450 cm⁻¹.

#### Investigation of Drug Loading in Albumin-Modified Liposomal Nanoparticles

In this section, to assess the amount of drug loaded within the liposomes, the supernatant of the centrifuged solution (the initial solution following liposome synthesis), which contained unencapsulated and free drug, was utilized. By examining the absorbance at the drug’s maximum wavelength and employing the linear equation derived from the standard curve, the amount of unencapsulated drug was determined. The solvent for the posaconazole liposomal nanoparticles, which is distilled water, was used as a blank in the spectrophotometer. The following equation was employed to calculate the drug loading percentage:

Loading=100×(Total Drug Amount−Free Drug Amount)Total Drug AmountL oading=100×Total Drug Amount(Total Drug Amount−Free Drug Amount) An evaluation of liposomal nanoparticles modified to encapsulate drugs is completed using this methodology.

#### Investigation of Drug Release from Liposomes

To evaluate the drug release from liposomes, it was first necessary to prepare a phosphate-buffered saline (PBS). For this preparation, 0.8 g of sodium chloride, 0.2 g of potassium chloride, 1.15 g of disodium phosphate, and 0.2 g of monopotassium phosphate were mixed and diluted to a final volume of 1000 milliliters. Concurrently, an appropriate amount of dialysis bag (with a cutoff of 12 KD) was taken and placed in distilled water at 100 degrees Celsius for 10 minutes to activate it. After activating the dialysis bag and preparing the PBS solution, 0.5 mg of posaconazole-loaded nanoparticles were dispersed in 4 milliliters of PBS and subsequently transferred into the dialysis bag, which was securely tied at both ends with dental floss. The dialysis bag was positioned at a 45-degree angle in a container holding 100 milliliters of PBS solution. At specified time intervals (1 hour, 2 hours, 4 hours, 6 hours, 8 hours, 12 hours, and then every 24 hours), 2 milliliters of the PBS solution were withdrawn as samples and replaced with fresh buffer. Finally, the absorbance of the collected samples was determined using a UV spectrophotometer, and by substituting the absorbance values into the standard concentration-absorbance curve, the concentration of the drug released from the nanoparticles was calculated. This article aims to provide a comprehensive approach to quantifying drug release kinetics in liposomal drug formulations so that a better understanding of liposomal drug formulations’ therapeutic efficacy and stability can be acquired.

#### *In vitro* evaluation of the efficacy of posaconazole-containing liposomes against mucormycosis-causing fungi

For this study, six fungal strains responsible for mucormycosis were utilized. All of these fungi were isolated from clinical samples of patients suffering from this disease and were accurately identified through genomic sequencing of the ITS1-5.8S rDNA-ITS2 region. To conduct antifungal tests, these isolates were sequentially numbered from 1 to 6, with number 1 designated as *Lichtheimia ramosa*, while the remaining numbers corresponded to *Rhizopus oryzae*, the most common causative agent of mucormycosis. The initial culture of the fungi was performed on Sabouraud dextrose agar. To prepare this culture medium, 65 g of the powder was dissolved in 1 liter of distilled water and boiled over a flame. The resulting solution was then autoclaved at 121 degrees Celsius and 1.5 atmospheres for 15 minutes to ensure sterility. After sterilization, the culture medium was poured into Petri dishes with a diameter of 6 centimeters and allowed to cool at room temperature. Following these steps, the prepared culture medium was transferred to a refrigerator for storage. Ultimately, as previously mentioned, this culture medium was used for the initial cultivation of the fungi, allowing clinical samples to incubate for 48 to 72 hours to achieve sufficient conidiation. This preparation is crucial for evaluating the antifungal efficacy of treatments against mucormycosis-causing fungi in vitro.

For the antifungal testing, the Clinical & Laboratory Standards Institute (CLSI) protocol, version M38, third edition, was utilized (20). Initially, RPMI 1640 culture medium with glutamine and without bicarbonate was prepared. To create the RPMI medium, 34.56 g of MOPS buffer was dissolved in 900 milliliters of distilled water, and the pH was adjusted to a range of 6.9 to 7.1 (if it was below this range, 1 N sodium hydroxide was used for adjustment). Finally, the solution volume was brought to 1 liter and thoroughly mixed. Subsequently, 10.4 g of culture medium powder was added to the buffered solution and stirred on a magnetic stirrer for 10 minutes until completely clear. Next, the resulting solution was filtered through a syringe filter and transferred into sterile Falcon tubes for storage in a refrigerator. After preparing the RPMI medium, a stock solution of posaconazole was also prepared for testing. According to the previously mentioned protocol, the required concentration of posaconazole for evaluating its antifungal effect against mucoralean fungi is 16 µg/ml; thus, a stock solution with a concentration 100 times higher (1600 µg/ml) was prepared. It is noteworthy that dimethyl sulfoxide (DMSO) was used as the solvent for preparing the stock solution. Similarly, a stock solution of liposomes containing posaconazole was prepared, with the only difference being that distilled water was used for dispersing the drug instead of DMSO. Simultaneously with the preparation of drug stocks, fungal suspensions were created for each isolate. For suspension preparation, three Falcon tubes with a volume of 15 milliliters were prepared for each isolate. The suspension process involved first using a sterile swab to collect conidia from the surface of cultured fungi from each isolate so that conidia adhered to the swab. Then, 5 milliliters of normal saline were added to one of the previously mentioned Falcon tubes, and the swab containing conidia was placed into this tube to ensure proper dispersion of conidia within the saline and create a suspension. After this step, the Falcon tube containing the initial suspension was allowed to stand undisturbed for 5 to 10 minutes so that larger fungal fragments settled at the bottom. Once the mycelial fragments settled, the upper portion of the suspension was gently transferred to a second Falcon tube in volumes of one milliliter until sedimentation was avoided. After transferring the fungal suspension to the second Falcon tube, its optical density was measured using a spectrophotometer at a wavelength of 530 nm. The acceptable absorbance range at this stage, according to CLSI protocol, was between 0.15 and 0.17. If the absorbance fell within this range, the fungal suspension was diluted at a ratio of 1:50 using RPMI in a third Falcon tube to prepare the final suspension. It is important to note that the final concentration of the prepared suspension ranged from 0.4×10^4^ to 5×10^4^ colony-forming units per milliliter. After preparing fungal suspensions for all isolates and creating drug stocks, a 96-well plate was utilized to evaluate the antifungal effects of both drugs. Each row of the plate corresponded to one of the isolates numbered from 1 to 6. In one column, 100 µl of fungal suspension and 100 µl of RPMI culture medium without any drug were added as a positive control; in another column, 200 µl of RPMI culture medium was added to ensure no contamination occurred in this medium. In other columns, 100 µl of fungal suspension was added along with serial dilutions of the drug (serial dilution performed using RPMI culture medium). The final drug concentrations in these columns were as follows: 16, 8, 4, 2, 1, 0.5, 0.25, 0.125, 0.0625, and 0.03125 µg/ml. It should be noted that one 96-well plate was used for posaconazole and another separate plate for evaluating liposomes containing posaconazole. After mixing the drug with the fungal suspension in the plate, it was incubated at 35 degrees Celsius for 24 hours, after which the results were read. Synthetic drugs and formulations can be used in vitro to assess antifungal activity against mucormycosis-causing fungi.

#### Fibroblast cell culture

The L929 mouse embryonic fibroblast cell line was maintained in F12 DMEM culture medium supplemented with 10% (v/v) fetal bovine serum (FBS) and 1% (v/v) antibiotics (penicillin/streptomycin) (complete medium) at 37 degrees Celsius in a humidified atmosphere with 5% carbon dioxide. The culture medium was replaced every 2 to 3 days, and when the cells reached approximately 70% confluency in the culture flask, they were passaged. This cultivation method ensures optimal growth conditions for the fibroblast cells, which are essential for subsequent cytotoxicity testing of the synthesized liposomal formulations.

#### Assessment of Cell Viability

The L929 cells were seeded in 96-well cell culture plates overnight to investigate cell viability. Subsequently, the drug posaconazole and posaconazole-containing liposomes, which had been sterilized using ultraviolet light, were added to each well at concentrations of 400, 200, 100, 50, and 25 µg/ml. The cells were then incubated for 24, 48, and 72 hours. It is important to note that these concentrations were selected based on previous studies that reported an IC50 of 175 µg/ml for posaconazole ((21)-

(22)). In each well, 100 µl of a 3-(4,5-dimethylthiazol-2-yl)-2,5- diphenyltetrazolium bromide (MTT) solution was added and incubated for an additional 4 hours after the incubation period. DMSO was added to each well and incubated for 20 minutes after all culture media had been removed.

Finally, cell proliferation was determined by measuring the absorbance of each well at wavelengths of 570 nm and 690 nm using a microplate reader. In this method, posaconazole and its liposomal formulations are evaluated for their ability to increase cell viability.

## Results

### Maximum Absorption Wavelength of Posaconazole and Its Standard Absorbance Curve

A spectrophotometer was used to measure posaconazole’s maximum absorption wavelength (max). Additionally, the standard absorbance curve for posaconazole is presented in Figure 1.

**Figure 1.**
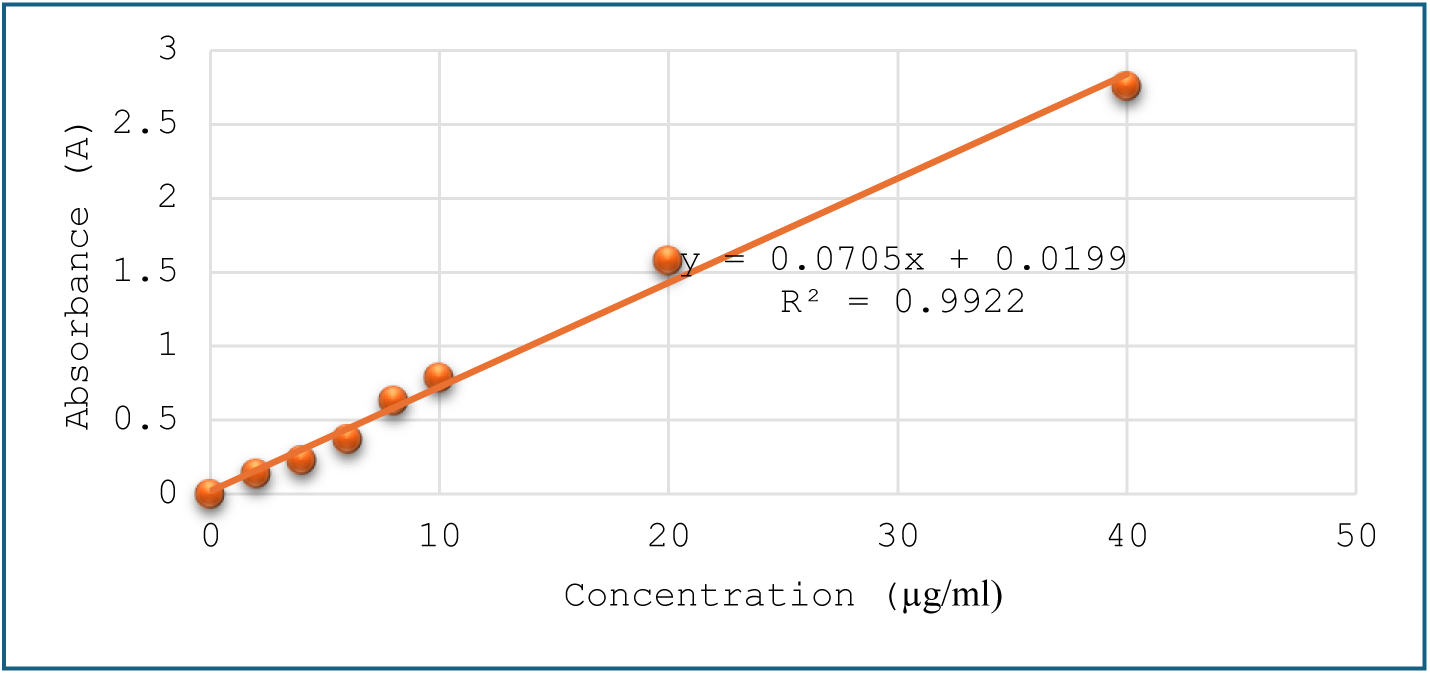
Standard Curve of Posaconazole at Different Concentrations.

### Results of SEM Imaging of Nanoparticles

The SEM images obtained from the nanoparticles are presented in Figure 2.

**Figure 2.**
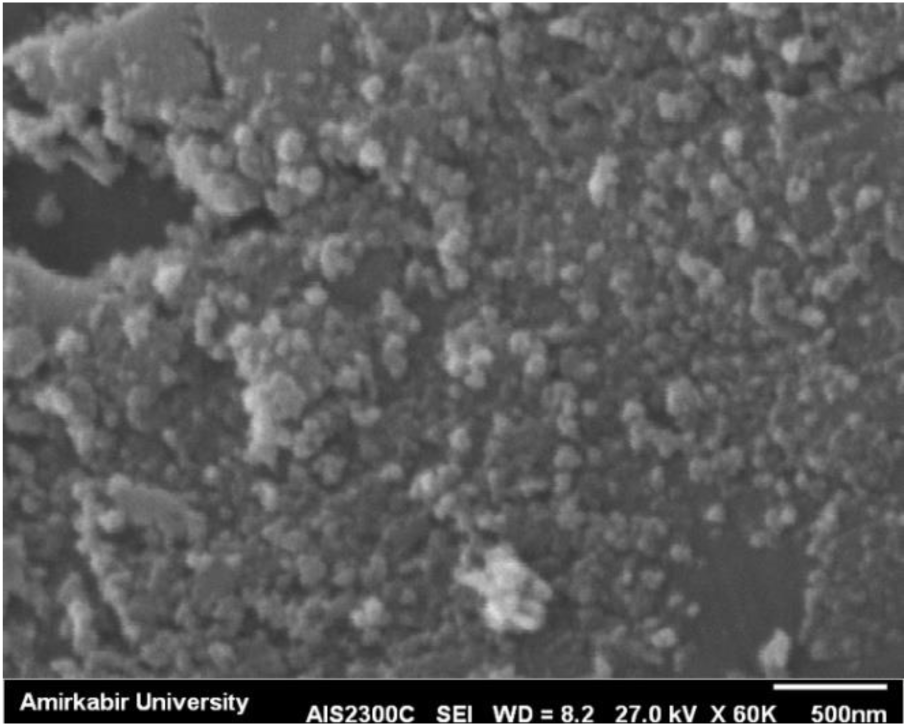
SEM Image of the Morphology and Shape of the Nanoparticles. Based on these images, the liposomes containing posaconazole have an approximate size of 60 nm.

### Results of DLS for Liposomal Nanoparticles

The DLS analysis of the synthesized posaconazole-loaded liposomal nanoparticles indicated that the size of these nanoparticles in their hydrated state was 126.6 ± 17.4 nm, with a polydispersity index of 0.7. Figure 3 illustrates the corresponding findings.

**Figure 3.**
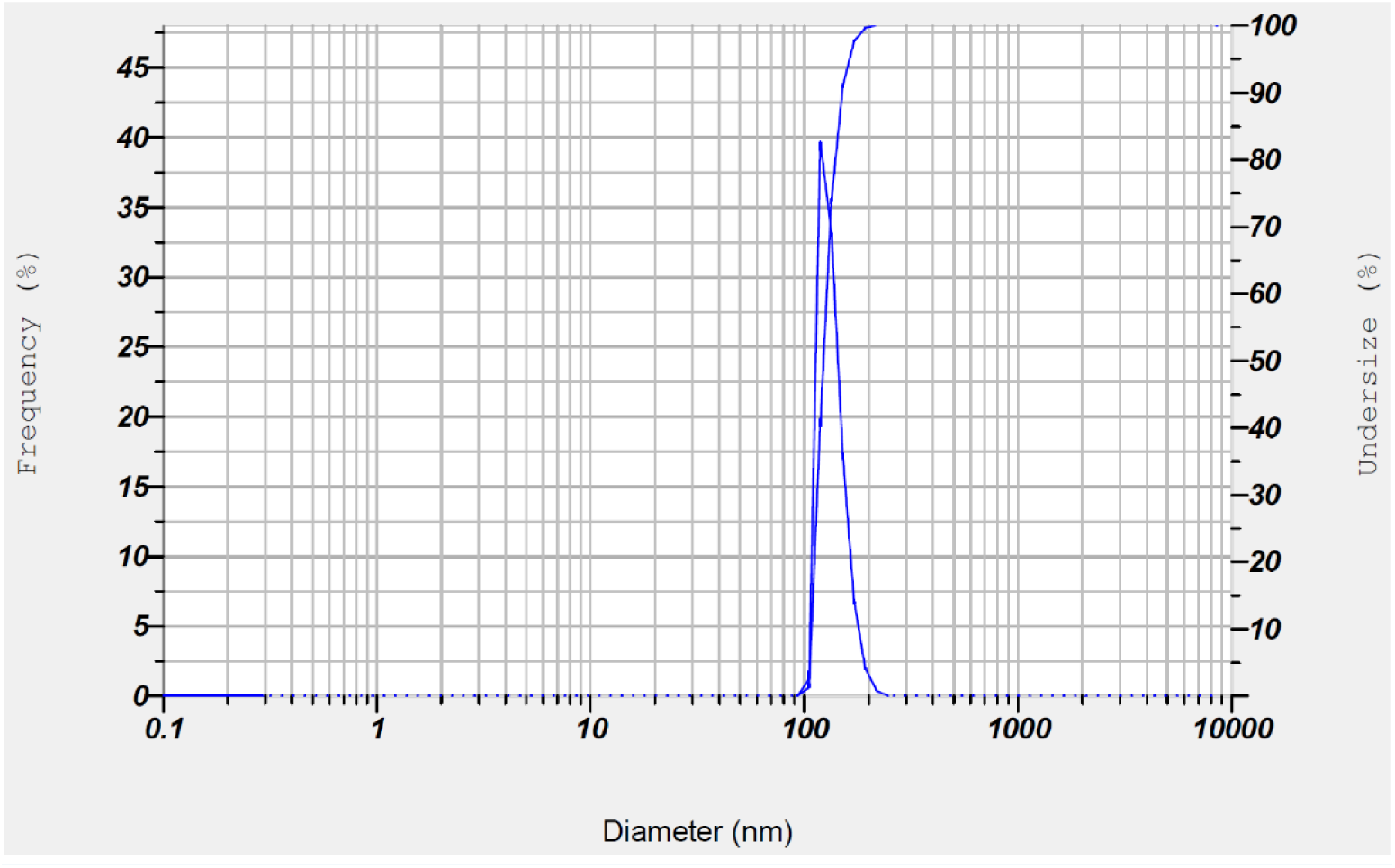
Results of Nanoparticle Size Analysis Using DLS. Additionally, in this test, a zeta potential value of -51 mV was obtained. Given that this value is less than -30 mV, it can be concluded that the synthesized nanoparticle exhibits adequate stability.

### Functional Group Analysis of Nanoparticles Using FTIR

FTIR analysis was employed to demonstrate the interactions between the lipid, posaconazole, and the chemical interactions among them in the synthesized nanoparticles. The results of the FTIR analysis are presented in Figure 4.

**Figure 4.**
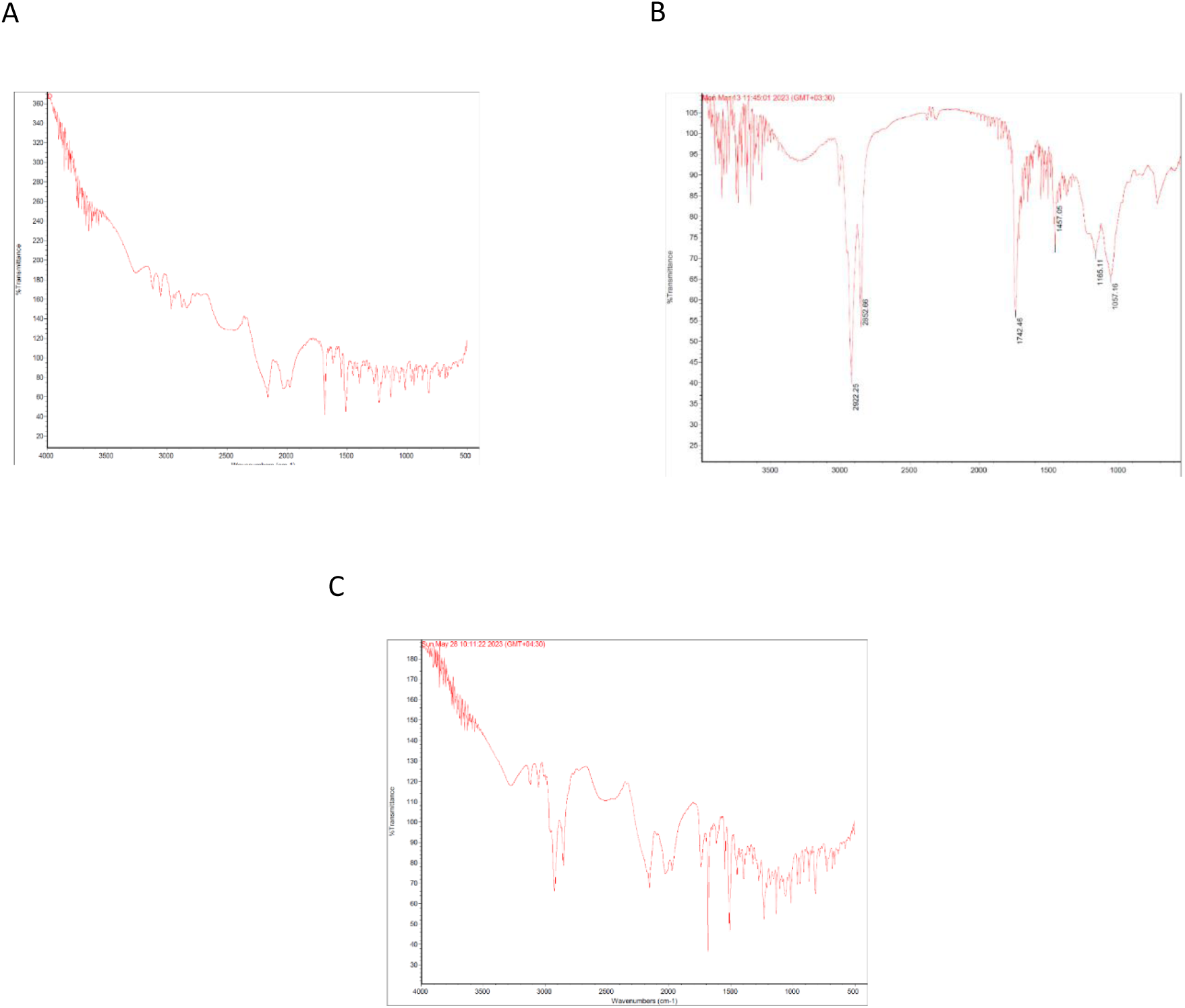
FTIR Spectra for A) Posaconazole B) Lipid C) Posaconazole-Containing Liposomes.

As shown in part A of the figure, the spectrum corresponding to posaconazole displays peaks that indicate various bonds within its structure. These peaks include those in the wavelength range of 2500 to 3000 cm^−1^, which represent C-H bonds; a peak at 1412 cm^−1^, indicating a furan ring; a peak at 1392 cm^−1^for alkyl C-H bonds; a peak at 1110 cm^−1^ for C-F bonds; and finally, peaks at 719 cm^−1^associated with aromatic C-H bonds. Part B of the figure illustrates the spectrum related to the lipid, where the peak at 2922 cm^−1^ corresponds to C-H bonds, and the peak at 1742 cm^−1^ indicates a C=O double bond. Finally, part C presents the FTIR spectrum of the synthesized nanoparticles, which also exhibits the peaks identified for both posaconazole and lipid. Therefore, based on these observations, it can be concluded that the nanoparticles have been successfully formed.

### Results of Posaconazole Loading in Nanoparticles

The supernatant solution was diluted 100-fold and analyzed as a control sample using distilled water in a spectrophotometer. The obtained absorbance value in this case was 0.096. Subsequently, the necessary calculations were performed to determine the percentage of drug loading.

0.096=0.0705x+0.01990.096=0.0705*x*+0.0199 x=1.07943262*x*=1.07943262

Concentration:

1.07943262×100=107.943262 g mL1.07943262×100=107.943262 g mL

Amount of free drug:

107.943262×20=2158.86524 g107.943262×20=2158.86524 g

Percentage of drug loading:

Drug loading percentage=(10000−2158.8652410000)×100Drug loading percenta ge=(1000010000−2158.86524)×100

Drug loading percentage=78.4113%Drug loading percentage=78.4113%

These results indicate that the drug loading efficiency of posaconazole in the nanoparticles is approximately 78.41%.

### Posaconazole Release from Nanoparticles in an In Vitro Environment

The results from the analysis of samples taken at specific time intervals indicated that over 70% of posaconazole is released into the medium within the first 24 hours. Following this initial release, the rate of drug release gradually decreases, such that by 120 hours (5 days), the drug is fully released into the environment. The release profile of posaconazole from the liposomal nanoparticles is illustrated in Figure 5.

**Figure 5.**
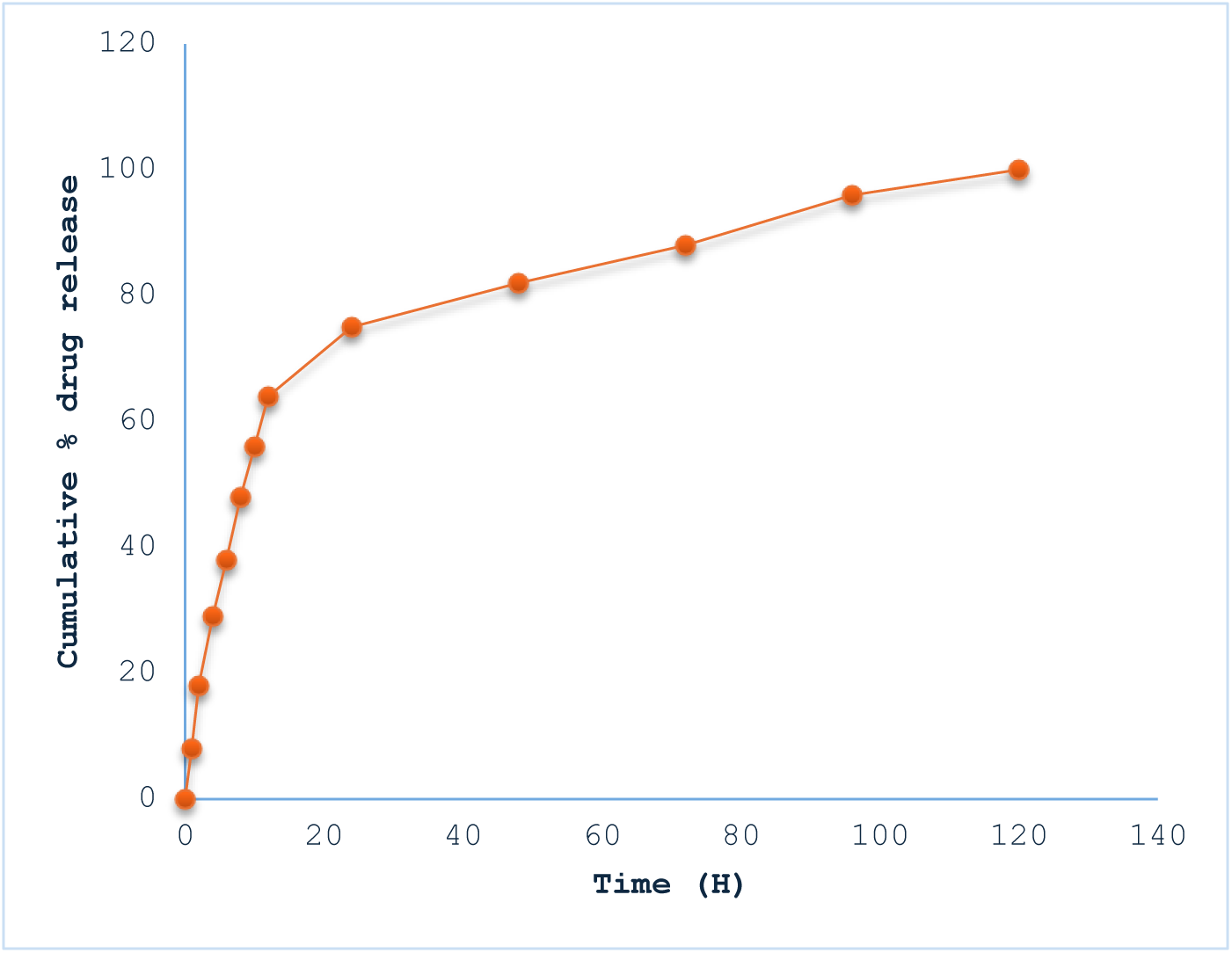
Release Profile of Posaconazole from Nanoparticles in an In Vitro Environment.

### Antifungal Activity of Synthesized Nanoparticles Against Mucormycosis-Causing Fungi in an In Vitro Environment

The results of the antifungal activity tests were read according to the previously mentioned CLSI protocol. The plates were visually evaluated, and the minimum concentration (µg/mL) of the drug at which 100% inhibition of fungal growth occurred— indicated by the absence of turbidity in the corresponding well—was considered the Minimum Inhibitory Concentration (MIC). Accordingly, Table 4-1 reports the MIC values for both posaconazole alone and posaconazole-loaded liposomes for each of the isolates numbered 1 to 6. As evident, the nanoparticles containing posaconazole exhibited MIC values that were either lower than or equal to those of posaconazole alone across all strains, indicating greater efficacy of the posaconazole liposomes against these fungi compared to the free drug. Additionally, Figure 6 displays the plate used for this assay.

**Table 1.** MIC Values for Posaconazole and Posaconazole-Loaded Nanoparticles for Each Isolate.

| **Drug**<br>**NO** | **Posaconazole** | **Posaconazole-**<br>**Loaded**<br>**Nanoparticles** |
| --- | --- | --- |
| **1** | 2 µg/ml | 1 µg/ml |
| **2** | 8 µg/ml | 2 µg/ml |
| **3** | 4 µg/ml | 2 µg/ml |
| **4** | 4 µg/ml | 2 µg/ml |
| **5** | 8 µg/ml | 2 µg/ml |
| **6** | 8 µg/ml | 4 µg/ml |

### Cytotoxicity Assessment of Posaconazole-Loaded Liposomes Using the MTT Assay

The cytotoxicity of the synthesized nanoparticles was evaluated by assessing the survival of normal fibroblast cells after exposure to the drug for 1, 3, and 5 days. These time points were chosen because, firstly, the therapeutic and antifungal effects of the drug are observed within the first 24 hours, and secondly, the entire drug is released from the nanoparticles over approximately 5 days. According to the results obtained, cell survival in all concentrations exceeded 70% during the first 24 hours, and no toxicity was observed at this time. It is noteworthy that at concentrations of 100, 200, and 400 µg/ml, although no toxicity was detected for either posaconazole alone or the posaconazole nanoparticles in the first 24 hours, the survival rate of cells exposed to liposomes was significantly higher than that of cells treated with posaconazole alone. In contrast to the first 24 hours, results obtained on day three indicated toxicity at a concentration of 400 µg/ml for posaconazole alone. Although posaconazole nanoparticles exhibited lower toxicity than posaconazole alone across all concentrations, these nanoparticles also resulted in less than 65% cell survival at a concentration of 400 µg/ml. Ultimately, results from the MTT assay on day five demonstrated that cell survival at concentrations of 25, 50, and 100 µg/ml for both posaconazole and posaconazole nanoparticles remained above 70%. In addition, while posaconazole nanoparticles showed lower toxicity at 400 µg/ml, cell survival fell below 60%. This indicates that the difference in toxicity between posaconazole nanoparticles and posaconazole- free had diminished. The results of this assay are illustrated in Figure 7.

**Figure 7.**
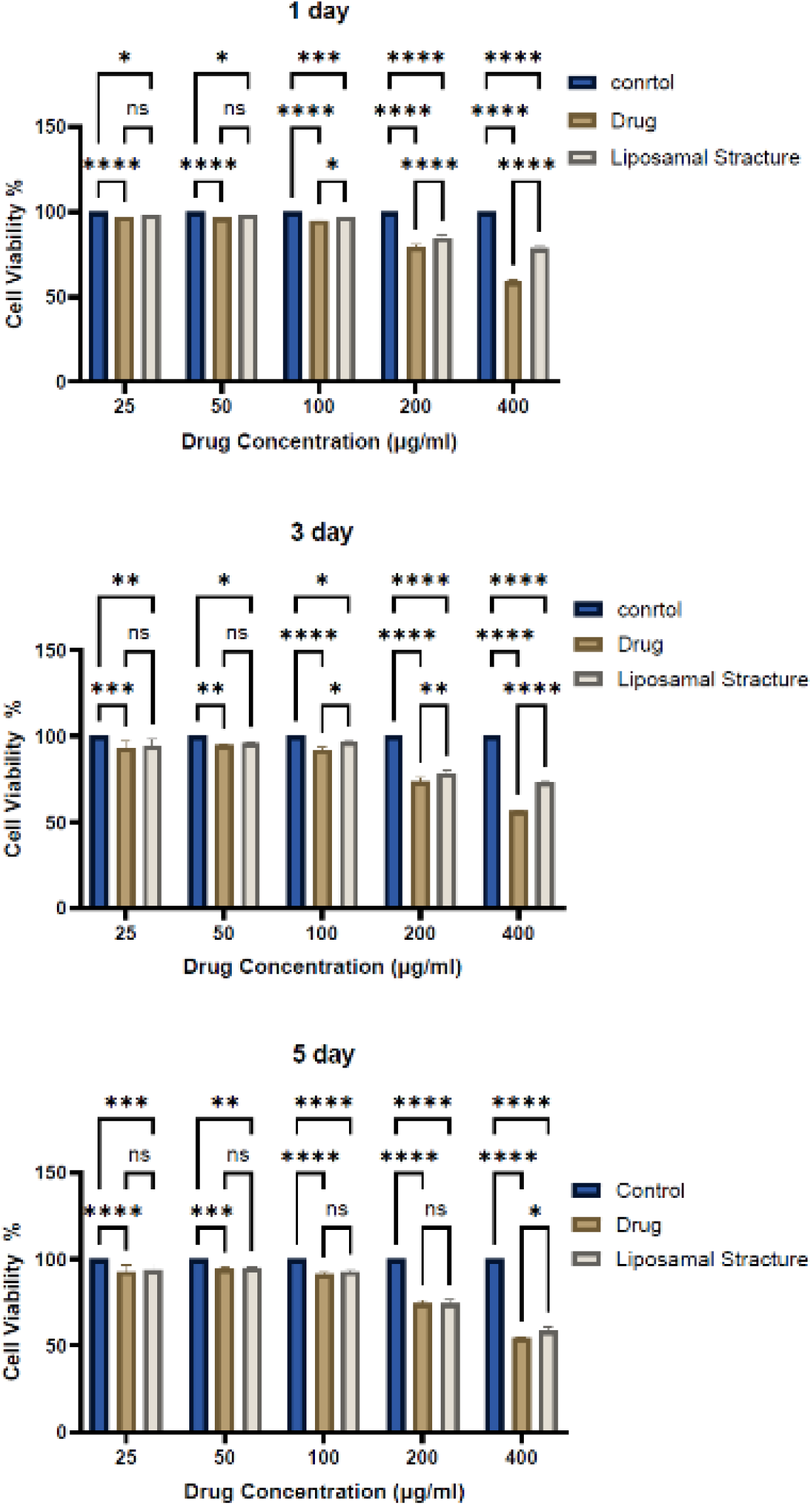
The result of the MTT test for nanoparticles of posaconazole and posaconazole alone.

Posaconazole, which received FDA approval in 2006, is recognized as one of the most effective and safe medications for treating a wide range of fungal diseases. In recent years, its application in the treatment of mucormycosis has also attracted attention. Although substantial evidence indicates the positive impact of posaconazole on mucormycosis, its use is associated with challenges such as a short half-life, low bioavailability, and consequently the need for frequent administration of high doses (23). In this context, various efforts have been made in recent years to provide suitable solutions to address these challenges. The use of novel drug delivery systems based on nanotechnology is one strategy currently proposed to resolve drug-related issues. Liposomes are one such drug delivery system that enhances efficacy, reduces toxicity, allows for sustained release, and increases the half-life of drugs (24). The present study aimed to overcome the challenges associated with posaconazole using this liposomal drug delivery system, and the results indicate that this goal has been achieved at least in vitro. In summary, liposomes were synthesized using the thin film layer method. The reason for choosing the thin film layer method over other liposome synthesis methods was its ease and availability compared to other techniques (19). To ensure posaconazole-containing liposome formation, FTIR testing was performed. This demonstrated lipid-drug interactions in the final sample, suggesting that the desired nanoparticles were correctly formed. In the next step, an ultrasonic probe device was utilized to control nanoparticle size in this study. After each synthesis stage, the resulting solution was sonicated with this device to minimize particle size as much as possible. Ultimately, based on DLS testing and SEM imaging, the morphology and size of the liposomes were examined. The results indicated that nanoparticles with an approximate size of 60 nm and a spherical shape were obtained, and the DLS scattering index was also acceptable. Additionally, the DLS device reported a zeta potential of -51 mv for the synthesized nanoparticles, which is reasonable given the negative charge of phosphatidylcholine and indicates complete stability of the nanoparticle structure. It is noteworthy that since this nanoparticle is intended for direct injection into the orbital cavity for treating rhino- orbit-cerebral mucormycosis, its 60-nanometer size is very suitable and will not cause any issues. This study examined the challenge of appropriate drug loading as one of the primary challenges in drug delivery systems. The results showed that the percentage of posaconazole loading within liposomes was approximately 80%. Furthermore, drug release from liposomal nanoparticles was measured over 5 days. Thus, these results suggest that using a liposomal form of posaconazole has resolved one of the challenges associated with this drug by increasing its half-life and prolonging its release over 5 days, therefore reducing the need for frequent administration; however, conclusive conclusions require further clinical and in vivo studies. These nanoparticles were evaluated and compared with posaconazole exclusively for their efficacy against mucormycosis-causing fungi, as well as their cytotoxicity. For antifungal activity assessment, CLSI protocol M38 was utilized, which is widely used in mycology to determine antifungal effects (20). The antifungal activity results indicated that the MIC of posaconazole nanoparticles against two species, Lichtheimia ramosa and Rhizopus oryzae—major causative agents of mucormycosis—were 1 microgram per milliliter and 2.29 µg/ml, respectively. In contrast, MIC values for posaconazole alone against these two species were 2 µg/ml and 6.06 µg/ml, respectively. Therefore, it is evident that the liposomal form effectively eliminates mucormycosis fungi at lower concentrations compared to posaconazole alone. Cell toxicity assessments conducted during the first 24 hours indicated that not only did posaconazole nanoparticles exhibit no toxicity, but they also showed lower toxicity at similar concentrations compared to posaconazole alone. However, results obtained on days three and five revealed that although liposomal posaconazole maintained lower toxicity than posaconazole alone, cell viability between these two began to converge. This observation confirms an increased half-life for the drug in its liposomal form since posaconazole alone diminished after 24 to 48 hours while the liposomal form continued releasing active drug and resulted in greater cell death. Based on cumulative results from antifungal testing and cytotoxicity assessments, it can be concluded that observed effects on fungi were not due to nanoparticle toxicity but rather therapeutic effects from the drug itself that led to fungal eradication; furthermore, the newly designed drug delivery system significantly enhanced drug efficacy. Despite previous uses of novel drug delivery systems for posaconazole, none have utilized the newly designed drug delivery system, which has significantly enhanced drug efficacy. Despite previous uses of novel drug delivery systems for posaconazole, none have utilized liposomes until now. In studies closest to this research, Osouli et al. (25) and Dourgan et al. (26) designed micelles containing posaconazole for ocular drug delivery. In Osouli’s study conducted in Iran in 2023, researchers employed thin film layer synthesis along with remote film loading and direct dispersion methods for micelle synthesis. They demonstrated that thin film layer synthesis yielded higher drug loading percentages with smaller nanoparticle sizes; thus, they chose this method for nanoparticle synthesis. Similarly, Dourgan’s study also used this same method for micelle synthesis. In both studies, different solvents and varying lipid-to-surfactant ratios were employed; changes in these parameters resulted in final nanoparticles taking a liposomal rather than a micellar form, unlike those observed in Osouli’s study. Comparing the liposomes from the current study with the final nanoparticles from Osouli’s and Dourgan’s studies reveals that their synthesized micelles had sizes of around 60 and 80 nm, respectively —similar to those produced here. In terms of drug loading percentages, Osouli reported around 80%, while Dourgan reported 90%, closely matching our findings of approximately 80%. However, concerning other assessed factors, liposomal nanoparticles exhibited superiority over micellar forms. Drug release from micellar forms was reported at only about 10 hours. Comparisons of antifungal activity between micellar forms and conventional posaconazole revealed no differences, according to Osouli’s study; however, Dourgan’s study reported stronger antifungal effects from micellar forms without providing MIC values for this effect. Notably, neither Osouli nor Dourgan assessed micellar toxicity. Overall, it can be concluded that liposomal nanoparticles perform better than the micellar forms designed by Osouli and Dourgan in terms of drug release duration and antifungal efficacy. Alongside micellar forms, lipid-based nanostructures (SLN) have also been employed for delivering posaconazole. In a study by Ghoreghaei et al.. (27), SLNs loaded with posaconazole were synthesized using high-speed homogenization methods. Although their drug loading results surpassed those found in this current study at around 96%, attributed to differences in synthesis methods and compounds used, their maximum drug release duration was only about 16 hours—far less than what has been observed with albumin-coated surface-modified liposomal posaconazole formulations. No assessments regarding toxicity or efficacy were conducted on synthesized nanostructures within their study. Therefore, an overall comparison between prior studies and current research indicates that utilizing liposomes as a novel drug delivery system for posaconazole is entirely new and exhibits more favorable effects compared to previously employed systems. In conclusion, based on the comparison of previous studies with the current research, it can be stated that the use of liposomes as a novel drug delivery system for posaconazole is entirely new and demonstrates more favorable effects compared to other drug delivery systems that have been utilized to date.

## Acknowledgements

We also express our gratitude to Sahar Soleimanzadeh for her dedication in this work.

